# Target identification of drug candidates with machine-learning algorithms: how choosing negative examples for training

**DOI:** 10.1101/2021.04.06.438561

**Authors:** Matthieu Najm, Chloé-Agathe Azencott, Benoit Playe, Véronique Stoven

## Abstract

(1) Background:Identification of hit molecules protein targets is essential in the drug discovery process. Target prediction with machine-learning algorithms can help accelerate this search, limiting the number of required experiments. However, Drug-Target Interactions databases used for training present high statistical bias, leading to a high number of false positive predicted targets, thus increasing time and cost of experimental validation campaigns. (2) Methods: To minimize the number of false positive predicted proteins, we propose a new scheme for choosing negative examples, so that each protein and each drug appears an equal number of times in positive and negative examples. We artificially reproduce the process of target identification for 3 particular drugs, and more globally for 200 approved drugs. (3) Results: For the detailed 3 drugs examples, and for the larger set of 200 drugs, training with the proposed scheme for the choice of negative examples improved target prediction results: the average number of false positive among the top ranked predicted targets decreased and overall the rank of the true targets was improved. (4) Conclusion: Our method enables to correct databases statistical bias and reduces the number of false positive predictions, and therefore the number of useless experiments potentially undertaken.

## 1. Introduction

Drug discovery often relies on the identification of a therapeutic target, usually a protein playing a role in a disease. Then, small molecular drugs that interact with the protein target to alter disease development are designed or searched for among large molecular databases. However, there has been a renewed interest in recent years for phenotypic drug discovery, which does not rely on prior knowledge of the target. In particular, the pharmaceutical industry has invested more efforts in rare diseases, which are often poorly understood, and for which no therapeutic targets may have been discovered. Although phenotypic drug discovery has made possible the identification of a few first-in class drugs [1], once a phenotypic hit is identified, not knowing its mechanism of action is a strong limitation to fill the gap between the hit and a drug that can reach the market [2]. More fundamentally, the target points at key biological pathways involved in the disease, helping to better understand its molecular basis.

The work we present here aims at helping identification of the protein targets for hit molecules from phenotypic screens. Identification of a drug target based solely on experiments is out of reach because it would require to design biological assays for all possible proteins. In that context, *in silico* approaches can reduce number of experimental tests by focusing on a limited number of high probable protein targets. Among them, Quantitative Structure-Activity Relationship (QSAR) methods were developed with that purpose [3]. They are efficient methods for the inverse problem of finding new molecules against a given target, when ligands are already known for this target. However, using them to identify the targets of a given molecule would require training a model for each protein across the protein space, which is not possible because many proteins have only few, or even no, known ligand.

Docking approaches can address this question [4], but they are restricted to proteins with known 3D structures, which is far from covering the human proteome.

In the present paper, we tackle target identification in the form of Drug-Target Interaction (DTI) prediction based on machine-learning (ML) chemogenomic algorithms [5]. They can be viewed as an attempt to complete a large matrix of binary interactions relating molecules to proteins (1 if the protein and molecule interact, 0 otherwise). This matrix is partially filled with known interactions reported in the literature and gathered in large databases such as the PubChem database at NCBI [6]. They can be used to train ML chemogenomic algorithms by formulating the problem as a binary classification task, where the goal would be to predict the probability for pairs (*m*, *p*) of molecules and proteins to interact. They can be used both to predict drugs against protein targets, or protein targets for a drug, the latter being relevant to our topic.

However, training a good ML chemogenomic model is hindered by biases in the DTI databases, such as whether the molecule for which one wishes to make predictions has known interactions or not [7]. An additional issue arises when the databases only contain positive examples of (*m*, *p*) pairs known to interact, but no negative examples of (*m*, *p*) known not to interact. In this context, it is classical to assume that most unlabeled interactions are negatives, and to randomly sample negative examples among them [8]. In this work, we explore how to best choose negative examples to correct databases statistical bias, and reduce the number of false positive predictions, which is essential to reduce the number of biological experiments required for validation of the true protein targets.

## 2. Materials and Methods

### 2.1. Datasets

ML algorithms for DTI predictions need to be trained on datasets of known DTIs in which proteins and molecules are similar to those for which predictions will be performed. Hit molecules in phenotypic screens for drug discovery are mostly drug-like molecules [9], and proteins will be human proteins. We used the DrugBank database (version 5.1.5) [10] to build our training dataset, because although much smaller than other databases like PubChem or ChEMBL, it provides high quality bio-activity information regarding approved and experimental drugs, including their targets, and contains around 17 000 curated drug-target interactions. Therefore, we built a dataset called DB-Database hereafter, that comprises all (*m*, *p*) DTIs reported in DrugBank involving a human protein and a small molecular drug. Overall, the DB-Database comprises 14 637 interactions between 2 670 human proteins and 5 070 drug-like molecules, which make up our positive DTIs. Because training a ML algorithm also requires negative examples, we added an equal number of negative DTIs to the DB-Database based on two methods:

- **Random sampling:** Negative examples were randomly chosen among the pairs (*m*, *p*) that are not labeled as a DTI but such that both *m* and *p* are in the DB-Database, under the assumption that most of unknown interactions are expected to be negative. This process was repeated 5 times, leading to 5 training datasets called RN-datasets (for Random Negatives-dataset) hereafter, differing only by their negative examples.
- **Balanced sampling:** To avoid biasing our algorithms towards proteins with many interactions, negative examples were randomly chosen among unknown DTIs, although in such a way that each protein and each drug appeared an equal number of times in positive and negative interactions. This process was also repeated 5 times, leading to 5 training datasets called RN-datasets (for Random Negatives-dataset) hereafter, differing only by their negative examples. Building this set of negative DTIs is not trivial, and we propose the following algorithm:

1. Each protein and molecule in the DB-Database has a counter corresponding initially to its number of known ligands or targets, respectively;
2. For each protein, starting from those with the highest counter to those with a counter equal to 1, molecules are randomly chosen among those not known to interact with this protein and whose counter is greater or equal to 1;
3. Each time a negative DTI is chosen, the counter of the corresponding protein and of the molecule is decreased by one unit;
4. The process is repeated until all proteins and molecules counters are equal to 0.

Overall, the RN-datasets and the BN-datasets share the same set of positive DTIs, which are those in the DB-Dataset, and their total number of negative DTIs are the same and equal to that of positive DTIs. All these datasets are provided as supplementary files.

Finally, to compare the performance of the algorithm trained on the RN-datasets or the BN-datasets when predicting targets for “difficult” molecules (hit molecules will be generally “difficult” molecules, i.e. with no or few known targets), we considered a small dataset of DTIs involving 200 drugs that have few known targets. This dataset built as follows: from the 5 070 molecules in the DB-Database, we kept approved drugs that do not have more than 4 targets. This lead to 560 drugs involved in 851 interactions among which we randomly selected 200 of these DTIs, involving 200 different drugs, defining the so-called 200-dataset, provided as supplementary file.

### 2.2. Machine-learning algorithm

We tackle the question of target identification by a chemogenomic approach that predicts DTIs with a Support Vector Machines (SVM) ML algorithm [11]. Briefly, the SVM is trained on a dataset of known DTIs and learns the optimal hyperplane that separates the (*m*, *p*) pairs that interact from those that don’t. Although SVM can use vector representations of the data (i.e. descriptors for proteins and molecules), thanks to the so-called “kernel trick” [12], they can also find this hyperplane based on particular similarity measures between (*m*, *p*) pairs of training dataset, and called kernel functions *K*, without requiring explicit representation of the data.

A general method to build a kernel on (*m*, *p*) pairs is to use the Kronecker product of molecule and protein kernels [13]. Given a molecule kernel *K_molecule_* and a protein kernel *K_protein_*, the Kronecker kernel *K_pair_* is defined by:

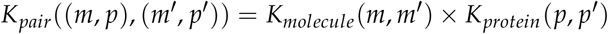

For proteins, we used a centered and normalized Local Alignment kernel (*LAkernel*), which mimics the Smith-Waterman alignment score between two proteins [14]. For the molecules, we used a centered and Tanimoto kernel, that uses molecular descriptors based on the number of fragments of a given length on the molecular graph [15].

The *LAkernel* has three hyperparameters: the penalties for opening (*o*) and extending (*e*) a gap, and the *β* parameter which controls the contribution of non-optimal local alignments to the final score. The Tanimoto kernel has one hyperparameter: the length *d* of the paths up to which paths on the molecule structure are considered. According to [8], we used the following values for these hyperparameters: *o* = 20, *e* = 1, and *β* = 1 for the *LAkernel*, and *d* = 14 for the Tanimoto kernel. The SVMs also requires a regularisation parameter classically called *C*, which controls the trade-off between maximising the margin (i.e. the distance separating the hyperplane and the two classes distributions) and minimizing classification error on the training points. This parameter was adjusted to *C* = 10 for both RN- and BN-datasets, based on a cross-validation scheme, as described in Section 2.3.

### 2.3. Training scheme and performance evaluation

We used a *K*-fold cross-validation (CV) scheme classically employed to train learning algorithms [16]:

1. Known positive DTIs of RN-datasets on the one side, and of negatives DTIS on the other side, were randomly split into *K* folds, and each negative fold was joined to a positive fold;
2. The model is run *K* times, each run using the union of (*K* − 1) folds as the training set, and the performance is estimated on the remaining fold;
3. Prediction performance are averaged over all folds.

More precisely, we first used a nested cross-validation scheme *Nested-CV* [17] to select the hyperparameter *C* of the SVM algorithm. It consists in a (*K* − 1) folds cross-validation *inner-CV* nested in a *K* folds cross-validation *outer-CV*. At each step of the outer CV, the *inner-CV* is repeated for the values of the *C* hyperparameter that were considered: 0.1, 1, 10, 100, 1000. The value providing the best CV prediction performance in terms of ROC-AUC or AUPR scores (see below) on average over all folds is retained. In our case, *C* = 10 leads to the best performances, both in terms of ROC-AUC and AUPR, on the RN-Dataset.

Finally, the outer K-fold cross-validation is performed, with *C* = 10. We used *K* = 5, a classical value in CV. This process was repeated 5 times for the 5 RN-datasets (see Section 2.1), and prediction performances were averaged over these 5 versions.

We estimated prediction performance using the following scores to judge the quality of the classifier:

- the Area Under the Receiver Operating Characteristic curve (ROC-AUC) [18]. The ROC curve plots true positive rate as a function of false positive rate, for all thresholds on the prediction score. Intuitively, the ROC-AUC score represents the probability that a positive interaction would be predicted by the classifier with a higher score than a negative interaction;
- the Area Under the Precision-Recall curve (AUPR) [19], which indicates how far the scores of true positives are from those of true negatives, on average;
- the Recall, representing the fraction of positive examples that are retrieved;
- the Precision, representing the fraction of true positives retrieved among predicted positives;
- the False Positive Rate (FPR), representing the fraction of true negatives among predicted positives.

The same scheme was employed when the algorithm was trained on the BN-Datasets, on which the same optimal value of *C* = 10 was obtained.

Finally, the algorithm is trained 5 times on the 5 complete RN-Datasets (or BN-Datasets) with *C* = 10, leading to 5 predictors. When scoring new DTIs, as for the 3 example drugs or for the 200-DB-Dataset (see Section 3.3 and Section 3.4), the algorithm provides 5 corresponding scores, and the final score is averaged over these 5 values.

## 3. Results

### 3.1. Performance of the SVM on the RN-datasets

In the present paper, we focus on identification of target candidates for phenotypic hit molecules based on a ML chemogenomic algorithm, which first requires to train the algorithm.

We considered a ML algorithm based on SVM, with the *LAkernel* [20] and the Tanimoto kernel [15] for proteins and molecules, respectively, because these methods displayed good prediction performances in previous chemogenomic studies, on average ([21],[22],[8]). The *LAkernel* is related to the Smith-Waterman score [14], but while the latter only keeps the contribution of the best local alignment between two sequences to quantify their similarity, the *LAkernel* sums up the contributions of all possible local alignments, which proved to be efficient for detecting remote homology.

Other kernels and other ML algorithms could have been used, such as Random Forests ([23],[24]), or Matrix Factorization ([25],[26]). However, our purpose was not to discuss the type ML algorithm or of kernels for target identification, but to study, whatever the ML algorithm might be, how to best train it for the particular task of target identification. More precisely, training a ML chemogenomic algorithm from a large DTIs database is in fact an example of Positive-Unlabelled (PU) learning problem because in practice, positive examples (known DTIs) are present in the training database, while non-positive interactions are in fact unknown interactions that have not been tested, or negative interactions that have not been published or included in database (most databases do not contain negative results). PU learning problems are usually converted into Positive-Negative (PN) learning problems for which many efficient ML algorithms are available. In chemogenomic, negative DTIs are classically randomly chosen among all unknown interactions, under the assumption that most of them are true negatives. Therefore, the algorithm was trained on the RN-dataset (for Random Negatives-dataset comprising positive and negative DTIs as defined in 2.1) based on a 5-fold cross-validation scheme as detailed in Section 2.2 and Section 2.3. A threshold of 0.5 in the probability score was chosen to discriminate between positive and negative predictions.

Table 1 shows the average performance of the SVM trained on the RN-datasets. The results in Table 1 show that overall, the SVM algorithm displays good performances, in the range of that provided by other state-of-the-art results obtained by other shallow or deep ML algorithms on a similar datasets ([8], [27]). In the context of target identification, it is important to limit the FPR, to avoid unnecessary experimental validation. However, a threshold of 0.5 over the probability scores was used to separate predicted positives from predicted negatives, as classically, although in practical cases, a higher threshold would be chosen to select target candidates, in order to reduce the number of experimental tests to the predictions with the highest confidence.

**Table 1.** Performances of DTI predictions of the SVM algorithm on the RN-datasets.

| AUPR | ROC-AUC | Recall | Precision | FPR |
| --- | --- | --- | --- | --- |
| $85.49 \pm 0.24$ | $88.03 \pm 0.13$ | $81.96 \pm 0.41$ | $93.3 \pm 0.43$ | $5.89 \pm 0.43$ |

We studied the distributions of probability scores for positive and random negative (presumably mainly true negatives) interactions, as predicted in the CV scheme. Figure 1 shows that these two distributions are well separated, and also suggests that a threshold around 0.7 in probability score could be used for high confident predicted positive interactions, in the RN-dataset. In addition, the rank of a predicted interaction is also an important criterion, because the goal of virtual screens is to drastically reduce the number of experiments to perform. In identification of hit molecules against a given therapeutic target, typically, the top 5% percent of the best ranked molecules are screened [28]. Usually, an experimental assay with a simple readout has been set up for the target of interest, which allows to evaluate relatively high numbers of candidate molecules selected in the virtual screen. The inverse problem of target identification is more difficult because validation requires to test the phenotypic hit molecule in a different biological assay for each predicted target considered for experimental evaluation. This obviously requires much time and efforts, because these assays may not all be available, and therefore, my have to be designed. This can be a real challenge if the function of a candidate target is not suitable to design a simple biological test. Therefore, we added the stringent but realistic threshold of top 1% in rank. In other words, in the following, we will consider positive predicted targets with scores above 0.7 and ranked in the top 1%, to simulate a realistic list of candidate targets that would be considered for experimental test. We discuss how to best train the algorithm in order to minimize the number of useless biological experiments that would be undertaken for false positive targets satisfying these two criteria, because this represents a real bottleneck for real-case studies. Consequently, in what follows, since the RN-dataset comprises 2 670 proteins, we will consider as candidate targets proteins with a probability score above 0.7 and rank smaller than or equal to 27.

**Figure 1.**
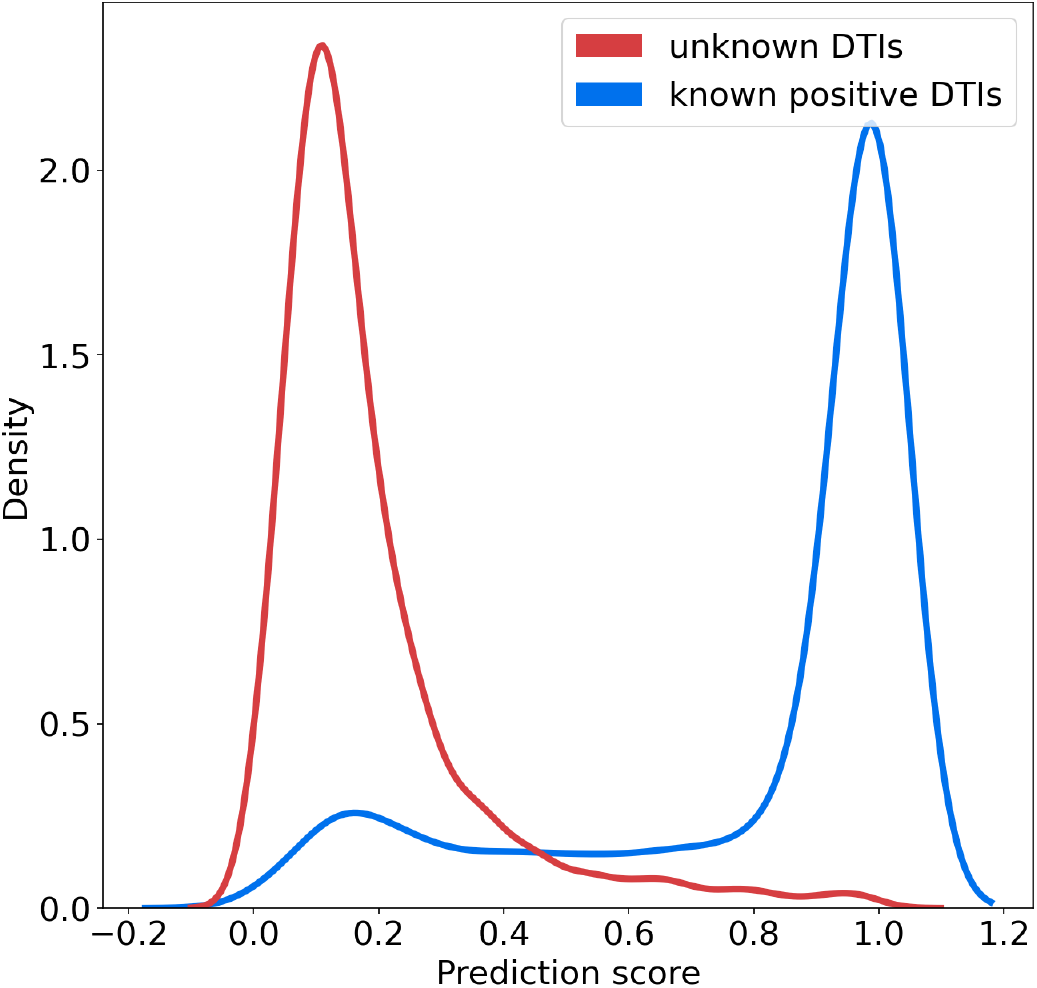
Distribution of the probability scores predicted for known positive DTIs and randomly chosen negative DTIs among unknown DTIs.

### 3.2. Statistical analysis of the DrugBank database

The DrugBank database [10] is a widely used bio-activity database. Although much smaller than PubChem or ChEMBL, it provides high-quality information for approved and experimental drugs along with their targets. It contains around 15,000 curated DTIs involving 2 670 human proteins (this set of proteins can be viewed as the “druggable” human proteome), and 5070 druglike molecules, corresponding to the DB-Database described in Section 2.1. This database is relevant for training of machine-learning models for DTI predictions involving human proteins and drug-like molecules. However, Figure 2 shows that there is a strong discrepancy between the number of known ligands per protein, or known protein targets per molecule.

**Figure 2.**
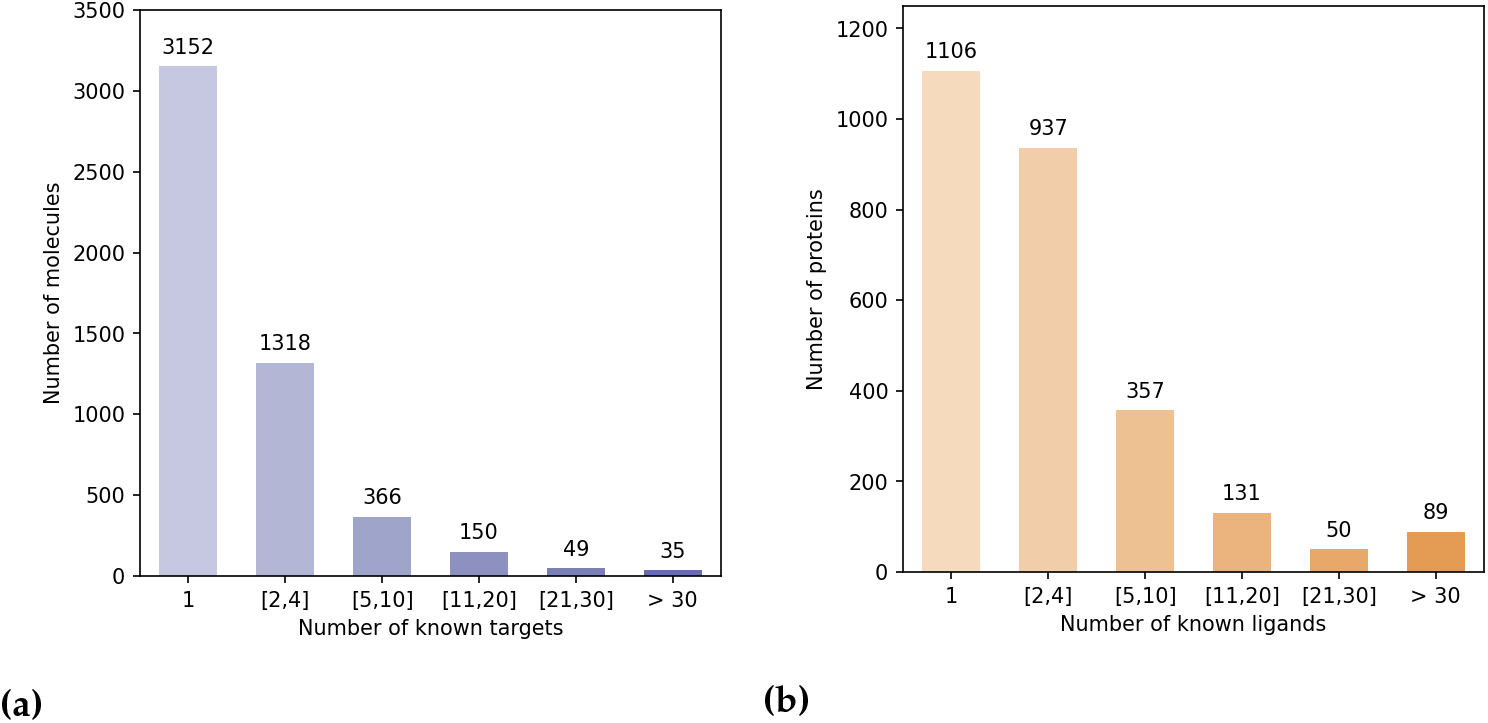
Statistical bias in the DB-Database. 2a Distribution of the molecules according to their number of targets in the DB-Database. 2b Distribution of the proteins according to their number of ligands in the DB-Database.

Indeed, the majority of proteins have 4 or fewer known ligands, while around 140 proteins have more than 21 ligands. We defined categories of proteins, depending on their number of known ligands (1, 2 to 4, 5 to 10, 11 to 20, 21 to 30, more than 30), and calculated the number of DTIs in the DB-database in each category. Overall, according to Table 2, 5.2% of the proteins are involved in 44% of DB-Database DTIs.

**Table 2.** Distribution in the DB-Database of the number of DTIs involving proteins from various categories, according to their number on known ligands.

| protein nb of ligands | nb of interactions |
| --- | --- |
| 1 | 1106 |
| 2 to 4 | 2527 |
| 5 to 10 | 2404 |
| 11 to 20 | 1920 |
| 21 to 30 | 1238 |
| > 30 | 5442 |

This bias arises from the fact that a few diseases like cancer or inflammatory diseases have attracted most research efforts, and many ligands have been identified against related therapeutic targets, compared to other less studied human proteins. For example, Prostaglandin G/H synthase 2, a well-known protein involved in inflammation, has 109 drugs reported at DrugBank. This statistical bias affects training of the SVM and is expected to perturb identification of targets for hit molecules, potentially by enriching top ranked proteins in false positive targets that have many known ligands.

### 3.3. Examples illustrating the impact of learning bias for target identification

Once trained, a ML algorithm identifies targets for a hit molecule by providing a list of proteins ranked by decreasing order of the estimated probability score all all (protein, hit) pairs. Candidate targets are chosen based on their probability score, their rank, and on potential prior biological knowledge that would highlight their relation to the considered disease. For example, a top ranked protein involved in cell cycle would be considered as a realistic candidate target for a hit identified in a cell proliferation screen in cancer research. The presence of many false positive targets among the top ranked proteins will not only lead to undertake useless experiments, but also potentially to discard true predicted targets pushed further down the list. Let us illustrate this problem in the case 3 molecules, randomly chosen among marketed drugs with only one known target in DrugBank. Assuming that their targets have been well characterized because they are marketed molecules, most other top ranked predicted targets will be false positive predictions. The 3 considered molecules are: alectinib (DrugBank ID DB11363, target: ALK), lasmiditan (DrugBank ID DB11732, target: HTR1F), and doxapram also known as angiotensin II (DrugBank ID DB11842, target: AGTR1). We orphanized these 3 molecules (i.e. we suppressed their single known target from the train set), as if they were hits from phenotypic screens, and used the SVM algorithm presented in Section 2.2 on the RN-dataset to predict their targets. For each molecule, the results consist in a list of the 2 670 proteins in the DB-Database, ranked by decreasing order of score.

As shown in the RN-dataset columns of Table 3, none of the known targets for those drugs are among the candidate targets as defined in Section 3.1. More precisely, for DB11363 and DB11842, although the probability scores of their known targets are above 0.7 (values 0f 0.8 and 0.76 respectively), their rank is 31 in both cases, above the the threshold of 27. For DB11732, the probability score of HTR1F is 0.67, with a rank of 107, and HTR1F would not either have been classified among the candidate targets for testing.

**Table 3.** DTI prediction results for 3 marketed drugs, when the algorithm is trained on the RN-Datasets or the BN-Datasets: number of False Positive predicted targets, score and rank of the true target

| drug | RN-datasets |  |  | BN-datasets |  |  |
| --- | --- | --- | --- | --- | --- | --- |
|  | FP | target score | target rank | FP | target score | target rank |
| DB11363 | 27 | 0.8 | 31 | 16 | 0.8 | 3 |
| DB11842 | 27 | 0.76 | 31 | 26 | 0.85 | 18 |
| DB11732 | 27 | 0.67 | 107 | 26 | 0.83 | 17 |

Analysis of the results highlighted that some of the best ranked candidate targets are frequent targets. For example, prothrombin F2 (120 ligands), cyclin dependant kisase CDK2 (137 ligands), and dopamine receptor 2 DRD2 (109 ligands) are top ranked predicted targets respectively for DB11842 (score of 0.97, rank 2), DB11732 (score 0.98, rank 1) and DB11363 (score 0.94, rank 5). The complete three ranked lists are provided in the Supplementary Materials.

These examples illustrate the impact of false positive predictions for target identification, because they can lead to discard even high-scoring true targets as for DB11363 and DB11842.

### 3.4. Choice of negative examples to correct statistical bias

Our observation that high-scoring false positives tend to have a large number of known ligands led us to make the assumption that the model trained using randomly sampled negative interactions is biased towards proteins with many known ligands, as well as possibly drugs with many known targets. This suggested us to choose negative DTIs in such a way that the training dataset contains, for each molecule and for each protein, as many positive than negative DTIs. The corresponding so-called BN-datasets (for Balanced Negatives-datasets) is detailed in Section 2.1. We recall that the BN-datasets contains the same positive DTIs as the RN-datasets, the former differing from the latter only by negative DTIs.

The SVM algorithm presented in Section2.2 was trained on the BN-datasets. As discussed above, for the problem of target identification, reducing the number of false positives among the top-ranked proteins is critical. Table 4 reports, for prediction score thresholds of 0.5 (usually considered) and 0.7 (considered in the present paper), the cross-validated FPR scores on these two training sets. It shows a strong statistical bias in FPR for the RN-datasets between proteins with few or with many known ligands, and that training on the BN-datasets greatly reduced this bias.

**Table 4.**
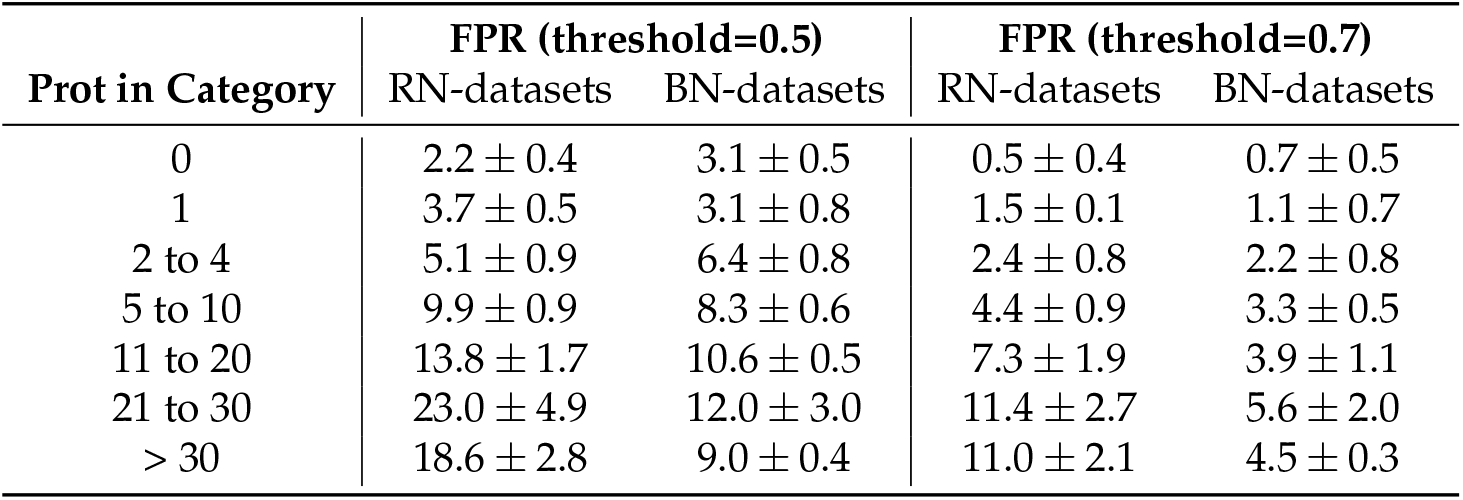
Rate of false positives for proteins with various numbers of known ligands.

| Prot in Category | FPR (threshold=0.5) |  | FPR (threshold=0.7) |  |
| --- | --- | --- | --- | --- |
|  | RN-datasets | BN-datasets | RN-datasets | BN-datasets |
| 0 | $2.2 \pm 0.4$ | $3.1 \pm 0.5$ | $0.5 \pm 0.4$ | $0.7 \pm 0.5$ |
| 1 | $3.7 \pm 0.5$ | $3.1 \pm 0.8$ | $1.5 \pm 0.1$ | $1.1 \pm 0.7$ |
| 2 to 4 | $5.1 \pm 0.9$ | $6.4 \pm 0.8$ | $2.4 \pm 0.8$ | $2.2 \pm 0.8$ |
| 5 to 10 | $9.9 \pm 0.9$ | $8.3 \pm 0.6$ | $4.4 \pm 0.9$ | $3.3 \pm 0.5$ |
| 11 to 20 | $13.8 \pm 1.7$ | $10.6 \pm 0.5$ | $7.3 \pm 1.9$ | $3.9 \pm 1.1$ |
| 21 to 30 | $23.0 \pm 4.9$ | $12.0 \pm 3.0$ | $11.4 \pm 2.7$ | $5.6 \pm 2.0$ |
| > 30 | $18.6 \pm 2.8$ | $9.0 \pm 0.4$ | $11.0 \pm 2.1$ | $4.5 \pm 0.3$ |

To highlight the impact of this bias correction in terms of target prediction, we show in Table 3 the prediction results for the 3 molecules discussed in Section 3.3, when the algorithm is trained with the RN-datasets or with the BN-datasets. When trained on the RN-datasets, none of the true targets would have been considered and a positive candidate target for testing, because of a score below 0.7 or a rank above 27, as discussed above. Training on the BN-datasets greatly improved the ranks and scores of the three true targets, and reduced the number of false positives, allowing the 3 corresponding true targets to fulfill the rank and score criteria defined in Section 3.1 to become candidate target for testing.

To better illustrate the interest of the proposed scheme for the choice of negative DTIs on a larger number of drugs we considered the 200-dataset consisting of 200 DTIs involving 200 marketed drugs with 4 of less known targets, as described in Section 2.1. This “difficult” test set was chosen because the aim was mimic newly identified phenotypic hits, for which known targets are expected to be scarce. For each drug, we artificially reproduced the process of target identification: the corresponding DTI was removed from the train set, a new SVM classifier was trained and used to score 2 670 DTIs involving this drug and all proteins of the DB-database. We compared the top-ranked predicted targets obtained when the algorithm is trained on the RN-datasets versus on the BN-datasets, as well as the number of removed false positive DTIs that would have been retrieved as candidates for testing (i.e. with a score above 0.7 and a rank lower or equal to 27).

Overall, training with the BN-datasets improved the predictions: the number of false positive DTIs decreased for 106 drugs, remained unchanged in 85 drugs, and increased in 9 drugs, as compared to training with the RN-datasets. In particular, this improvement allowed one additional true positive interaction to reach a score above 0.7 and a rank below 27 : 104 true targets were retrieved as candidates when training with BN-datasets, compared to 103 when training with RN-datasets. For the corresponding 104 drugs, the number of false positives decreased by 2.9 in average, and the rank of the true interactions decreased by 1.8 in average, bringing them even closer to the top ranked predicted proteins, and more likely to be chosen for experimental validation. Consistent with the results in Section 3.3 for the 3 example molecules, on average over the 200 considered molecules, the number of useless experiments potentially undertaken would have decreased when training with the BN-datasets.

## 4. Discussion

The present study highlights that despite overall good performances of ML algorithms in chemogenomics, as evaluated by cross-validation procedures, the identification of true positive protein targets for phenotypic hit molecules in real case studies may become a challenge when the algorithm is trained on a biased dataset. In such studies, biological experiments are guided by the predicted scores and ranks of candidate targets. Training on a biased dataset may lead not only to conduct useless experiments, but also to discard true positive targets because their scores are below the considered threshold, or because their rank is too high due to the presence of false positives among the top-ranked proteins. This point is rarely discussed in ML chemogenomic studies, which usually focus on cross-validation schemes that does not correspond to real case applications. We showed that choosing an equal number of positive and negative DTIs per molecule and per protein helps decrease the FPR in biased datasets. In the present study, we used a DrugBank-based dataset, despite its strong bias in terms of number of protein targets per molecule, or of ligand molecules per protein, because it is a relevant choice to train an algorithm for predicting interactions between drug-like molecules and human proteins.

Although we are aware that other and larger DTIs databases could have been used, the purpose of our study was not to discuss the choice of training set, in particular because other databases will also present the same type of bias as the DrugBank, for the same reasons. We also chose not to discuss the choice of the ML algorithm, since the SVM algorithm used here displays overall state-of-the-art performances, and since other ML algorithms can also be expected to suffer from the statistical bias present in the learning dataset. In fact, we recommend to balance the number of positive and negative DTIs per molecule and per protein for the training of any machine learning algorithm on any database.

To illustrate the advantage of the proposed scheme for the choice of negative interactions, we chose a threshold of 0.7 over the probability scores to identify candidate targets for experimental testing, although proteins with scores above 0.5 are classified as positives. This threshold of 0.7 was guided by the results in Figure 1, in order to select highly probable positive targets. It can be adjusted to a different value if the algorithm is trained with other databases, whether through the same kind of plot, or through a ROC-curve to correspond to a predefine false positive rate.

We added the stringent threshold of top 1% in rank for positive targets that would be tested, and this threshold could also be changed depending on available resources for experimental validation. However, the issue we identified and addressed in this paper does not depend on the scores and rank thresholds used, and choosing equal numbers of positive and negative DTIs per molecule and per protein for the training set will limit the number of false positives independently of the choice these thresholds. Finally, although the proposed scheme for the choice of negative examples was presented here in the context of target identification for hit molecules, it is of general interest and would be applicable to other types of PU learning problems when bias is present in the training set, which is a very common situation, in particular in many biological databases.

## Author Contributions

Conceptualization, M.N, V.S. and C-A.A.; methodology, M.N, C-A;A and V.S.; software, M.N and B.P.; validation, M.N; formal analysis, M.N and V.S.; investigation, M.N.; resources, M.N.; data curation, M.N; writing—original draft preparation, V.S and M.N; writing—review and editing,M.N., V.S., and C-A.A.; visualization, M.N.; supervision, V.S.; project administration, V.S; funding acquisition, V.S.

## Data Availability Statement

Datasets and results, presented in this study, are available at https://github.com/njmmatthieu/dti_negative_examples_data.git, included a README.md file describing them.

## Abbreviations

The following abbreviations are used in this manuscript:

BN-dataset: Balanced Negatives-dataset
RN-dataset: Random Negatives-dataset
DB-Database: DrugBank Database
DTI: Drug-Target Interaction
(m, p): (molecule, protein)
ML: Machine-Learning
SVM: Support Vector Machine (algorithm)

